# Plant infection by the necrotrophic fungus *Botrytis* requires actin-dependent generation of high invasive turgor pressure

**DOI:** 10.1101/2023.12.15.571920

**Authors:** Tobias Müller, Jochem Bronkhorst, Jonas Müller, Nassim Safari, Matthias Hahn, Joris Sprakel, David Scheuring

## Abstract

The devastating pathogen *Botrytis cinerea* infects a broad spectrum of host plants, causing great socio-economic losses. The necrotrophic fungus rapidly kills plant cells, nourishing their walls and cellular contents. To this end, necrotrophs secretes a cocktail of cell wall degrading enzymes, phytotoxic proteins and metabolites. Additionally, many fungi produce specialized invasion organs that generate high invasive pressures to force their way into the plant cell. However, for most necrotrophs, including *Botrytis*, the biomechanics of penetration and its contribution to virulence are poorly understood. Here we use a combination of quantitative micromechanical imaging and CRISPR-Cas guided mutagenesis to show that *Botrytis* uses substantial invasive pressure, in combination with strong surface adherence, for penetration. We found that the fungus establishes a unique mechanical geometry of penetration that develops over time during penetration events, and which is actin cytoskeleton dependent. Furthermore, interference of force generation by blocking actin polymerization was found to decrease *Botrytis* virulence, indicating that also for necrotrophs, mechanical pressure is important in host colonization. Our results demonstrate for the first time mechanistically how a necrotrophic fungus such as *Botrytis* employs this “brute force” approach, in addition to the secretion of lytic proteins and phytotoxic metabolites, to overcome plant host resistance.

## Introduction

Plant pathogens need to overcome preformed host barriers such as the cuticle and the cell wall for successful host invasion. Unless natural openings such as stomata or wounding can be exploited, pathogens have to create penetration sites. To this end, they can use polymer degrading enzymes, physical force or a combination of both^1^. The necrotrophic fungus *Botrytis cinerea* (*Botrytis* hereafter) belongs to the economically most devastating pathogens, causing gray mold on numerous vegetables, fruits and other crop plants^2^. Successful infection of necrotrophs is characterized by rapid killing of plant cells and depend on the secretion of cutin and cell wall degrading enzymes, and cytolytic and phytotoxic proteins^3^. While these proteins act redundantly and together are crucial for *Botrytis* virulence^4^, the role of physical force for infection success is poorly understood.

In contrast to necrotrophs, biotrophic pathogens secrete very few lytic enzymes and usually rely on the establishment of specialized infection structures^5,6^ to invade their host cells, causing only limited damage^7^. Infection structures include appressoria, penetration hyphae, infection hyphae and haustoria which grow inside plant tissue. Originally, melanin impregnation of the appressorium wall was believed to be critical to generate forces high enough to penetrate plant cell walls. The highly specialized appressorium of *Magnaporthe oryzae*, and to a lesser extent *Colletotrichum graminicola*, allow the generation of remarkably high maximal turgor pressures. For mature *M*. grisea a turgor pressure of 80 bar has been estimated^8^, and optical measurements of indentations on artificial substrates by *C. graminicola* revealed pressures up to 53.5 bar^9^. The belief that such force-based penetration is dependent on melanin-fortified appressoria has been challenged by the observation of an invasive force of 51.3 bar exerted by appressoria of the Asian soybean rust *Phakopsora pachyrhizi* which do not contain melanin^10^. Recently, pressures up to 28 bar were measured in the oomycete *Phytophthora infestans* even without the formation of an visible appressorium^11^.

In general, infection structures of necrotrophs are less specialized than those of biotrophs. The *Botrytis* appressorium lacks distinct morphological features and consists mainly of a terminal thickening of the hyphal tip. However, already in 1895 it was described that hyphae of *Botrytis* were able to penetrate gold-coated foils^12^; as these are inert to the secreted lytic cocktail, this implies that also this nectrotroph is capable of generating significant invasive pressures. To our knowledge, however, neither a quantitative description of turgor pressure generation nor a description of penetration mechanics has been reported for *Botrytis* or other necrotrophs.

Here we show that the necrotrophic fungus *Botrytis* employs unique spatio-temporal penetration mechanics. During subsequent stages of penetration, fungal hyphae are raising above incipient appressoria leading to an almost perpendicular entrance angle. In conjunction, the geometry of the adhesive site, that glues the appressorium to the host surface, changes from polarised to nearly isotropic. Moreover, measurements of the penetration kinetics and turgor pressure reveal that the invasive pressure grows up to 5 bar, while the maximally generated turgor pressure is more than 40 bar. Finally, we provide evidence that generation of turgor pressure is dependent on the actin skeleton and is required for full virulence of the necrotroph *Botrytis*.

## Results

To determine the biomechanical behaviour of *Botrytis* during penetration, we used fluorescently-labelled polydimethylsiloxane (PDMS) in combination with confocal laser-scanning microscopy. PDMS represents an artificial substrate resembling in stiffness and hydrophobicity a plant leaf^11^, and confocal microscopy allows to follow penetration in depth (z-direction). Initially, *Botrytis* conidia were inoculated on PDMS-coated coverslips and penetration was observed after 12h of incubation. We found that *Botrytis* development on PDMS occurs in a manner similar to that on glass (Supplementary Fig. 1), involving germination of conidia, appressorium formation and initiation of penetration. Maximum projections of acquired z-stacks reveal a clear displacement of fluorescently-labelled PDMS under the appressoria, indicating successful penetration (Fig. 1a). Notably, thinner hyphae emerge from appressoria, resembling penetration hyphae (Supplementary Fig. 2). For further investigation, we used the previously established fluorescent *Botrytis* strain *Bcgfp1*^13^ which allowed generation of 3D-reconstructions (Fig. 1b). Comparison of wildtype *Botrytis* and *Bcgfp1* revealed similar plant infection behaviour, indicating that GFP-expression does not affect virulence (Supplementary Fig. 3).

**Fig. 1:**
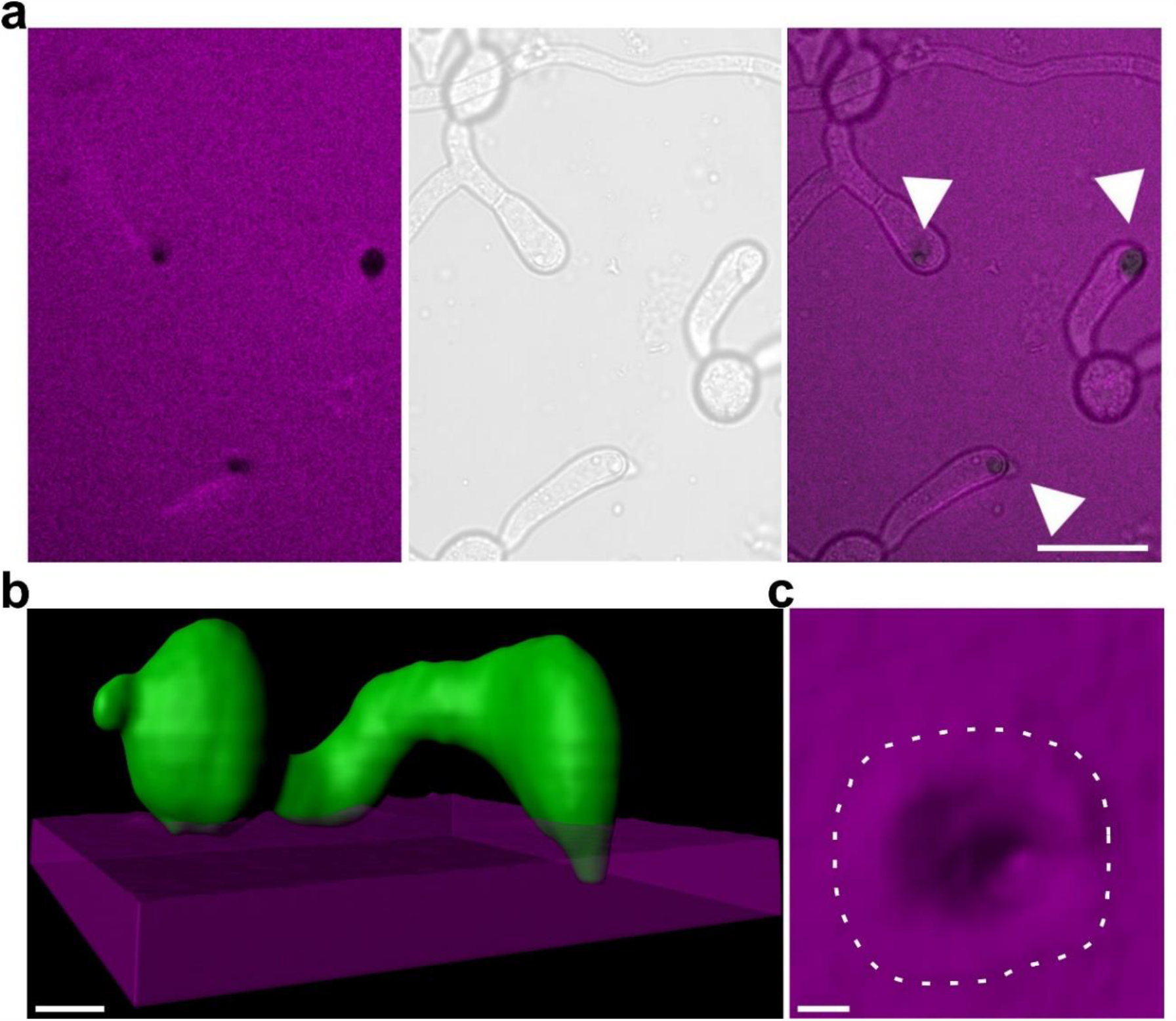
Development of *Botrytis* conidia on fluorescently-labelled PDMS. **(a)** Maximum projection, bright field and merged image of *Botrytis* on PDMS-coated coverslips. Arrowheads indicate appressoria. Scale bar is 10 μm. **(b)** 3D model (Imaris) of spores from the GFP-expressing strain *Bcgfp1* after 12h. Scale bar is 3μm. **(c)** 3D model of surface indentation. Scale bar is 1 μm.

To follow progressive stages of penetration, we recorded z-stacks after defined time points and rendered representative 3D models (Fig. 2a-d). Here, we could observe a unique pattern of penetration: Beginning around 10h after inoculation, indentations of PDMS were observed, and the hyphal tip was penetrating in an angle which changed during the penetration process. It was initially reminiscent of the so-called *naifu* penetration (Fig. 2a) observed for *Phytophthora*^11^ but changed later on (Fig. 2b). After 14h, and more clearly after 18h, the penetration angle became more perpendicular in parallel with deeper indentations (Fig. 2c, d). Furthermore, we observed increased arching of the hypha close to the appressorium resulting in a visible hunch after 18h (Fig. 2d). Based on the generated 3D models we estimated the penetration angles during different stages. Quantification revealed a significant shift from around 42° to 67° (Fig. 2e, f).

**Fig. 2:**
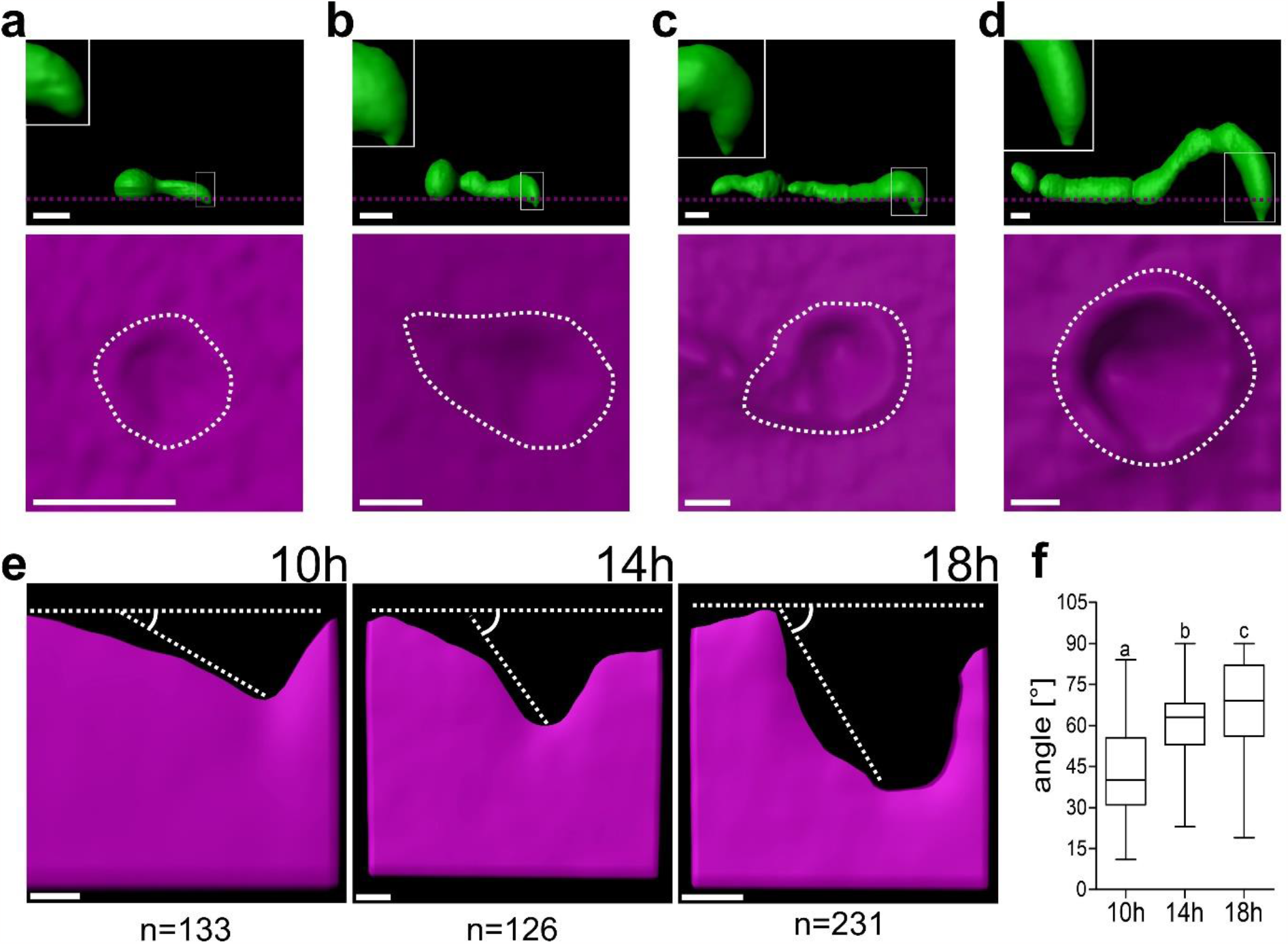
Kinetics of *Botrytis* penetration reveals a unique pattern. **(a-d)** Penetration kinetics of fluorescent *Botrytis Bcgfp1* on PDMS-coated coverslips after 6h (a), 10h (b), 14h (c) and 18h (d). Top part shows side view of elevation of a penetrating spore, the inserts showing close-ups of the penetrating tip. Scale bars are 5 μm (a, b, c) and 3 μm (d). Below (in magenta) indentations of PDMS are shown. Scale bars are 1 μm (a) and 3 μm (b, c, d). **(e)** Penetration angles after 10h (n=133), 14h (n=126) and 18h (n=231). Dotted lines show the measured angle between surface and the tip of the penetration hypha. Scale bars are 2 μm. **(f)** Box plot showing the measured angles at each time point. Box limits represent 25th-75th percentile, the horizontal line the median and whiskers minimum to maximum values. Statistical differences were analyzed by One-Way ANOVA with Tukey post hoc test, letters indicating significant differences; p<0,01. Experiments were repeated twice with similar results; representative models are shown.

Next, we quantified the substrate deformations induced by the indentation pressure of the *Botrytis* hypha. To this end, z-stacks that spanned approximately 20 μm in the z-direction and an axial resolution of 0.5 μm were generated starting at the boundary layer of PMDS and air (Fig. 3a). To capture conidia contact sites and to follow penetration, 20 z-slices were recorded above and below the boundary point (Fig. 3a). From the generated stacks we measured the intensity profile across the substrate–medium interface in each pixel. Fitting these intensity profiles to a sigmoidal function, we could extract surface height Δ*h* (Supplementary Fig. 4) to display surface-deformation maps (Fig 3b, c). In general, we observed zones of adherence (upward deformations) around the hyphae and indentation (downward deformations) beneath the centre of the appressoria. Earlier time points display a polarized mechanical geometry (Fig. 3b) which is followed by a more concentric circular symmetry later (Fig. 3c). Zones of adherence were positive for concanavalin A (a lectin binding glucosyl and manosyl groups), indicating that carbohydrates binding allows *Botrytis* attaching to the (artificial) surface (Supplementary Fig. 5).

**Fig. 3:**
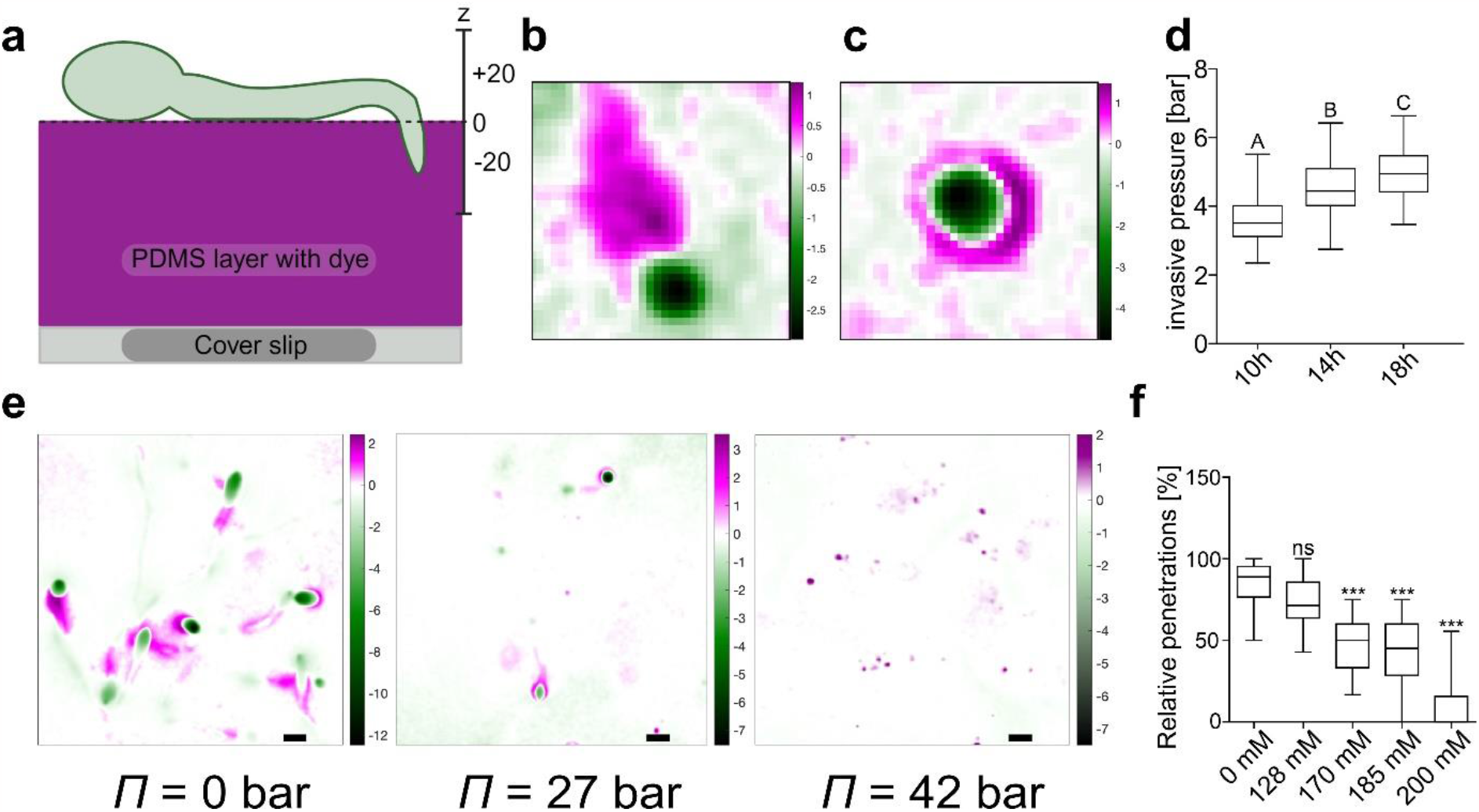
Surface deformation maps to calculate invasive and maximum turgor. **(a)** Model of the setup used to create surface-deformation map. **(b and C)** Surface deformation maps after 10h (b) and 18h (c). **(d)** Invasive pressure at 10h (n=50), 14h (n=61) and 18h (n=51). Significant differences were analyzed by One-Way ANOVA with Tukey post hoc test. Different letters indicate significant differences; p<0,01. **(e)** Surface deformation maps of *Botrytis* on PDMS generating an osmotic counter pressure via addition of different PEG2000 concentrations. **(f)** Box plot showing the relative amount of penetrations using 0 mM (n=288), 85 mM (n=28), 128 mM (n=207), 170 mM (n=102) 185 mM (n=112) and 200 mM (n=55) PEG2000. Box limits in the graph represent 25th-75th percentile, the horizontal line the median and whiskers minimum to maximum values. Significant differences were analyzed by One-Way ANOVA with Dunnet post hoc test, ***p<0,001; ns=not significant. Experiments were repeated twice with similar results; representative surface deformation maps are shown.

In walled cells, the invasive pressure, i.e. the pressure applied by the pathogen at the penetration site, is generated by the focussing of the internal cellular turgor pressure to a distinct site. To calculate the invasive pressure, we extracted from the surface deformation maps (Fig. 3) for each penetration site both the depth and width of the indentation site. Assuming a Hertzian contact law, and having previously determined the stiffness of the artificial substrate^11^, this allows us to calculate the indentation pressure generated by the pathogen at the invasion site. After 10h we found an invasive pressure of 3.7 bar which was steadily increasing during penetration until it reached 5.0 bar after 18h (Fig. 3d). For *Phytophthora*, it has been shown that the local invasive pressure accounts only for approximately 10-20% of the maximal cellular turgor^11^. To determine the maximum turgor pressure, we increased the osmotic pressure Π of the medium until suppression of penetration by *Botrytis* was almost complete^14^ (Fig. 3e). This was only reached at osmotic pressures between 27 bar and 42 bar^15^ confirming that also for *Botrytis* only 10-20% of the maximum turgor pressure is converted into locally applied, invasive pressure (Fig. 3f).

Since it has been reported that rearrangements of the actin cytoskeleton are a prerequisite for host penetration of *Phytophthora*^16^, we asked if actin has a similar role during *Botrytis* penetration. To interfere with actin filament polymerization, we used the toxin Latrunculin B (LatB) at 500 nM and 1 μM concentration. Latrunculin has been shown to block actin polymerization in most eukaryotic organisms including fungi^17,18^. Germination was unaffected within this range (Supplementary Fig. 6) but the penetration frequency and depth was suppressed visibly starting with 500 nM LatB (Fig. 4a). In addition, we observed structural changes of appressoria when treated with LatB (Fig. 4a, insert). This is somewhat reminiscent of hyphal blunting described for Latrunculin-treated *Phytophthora* during *naifu* invasion^16^.

**Fig. 4:**
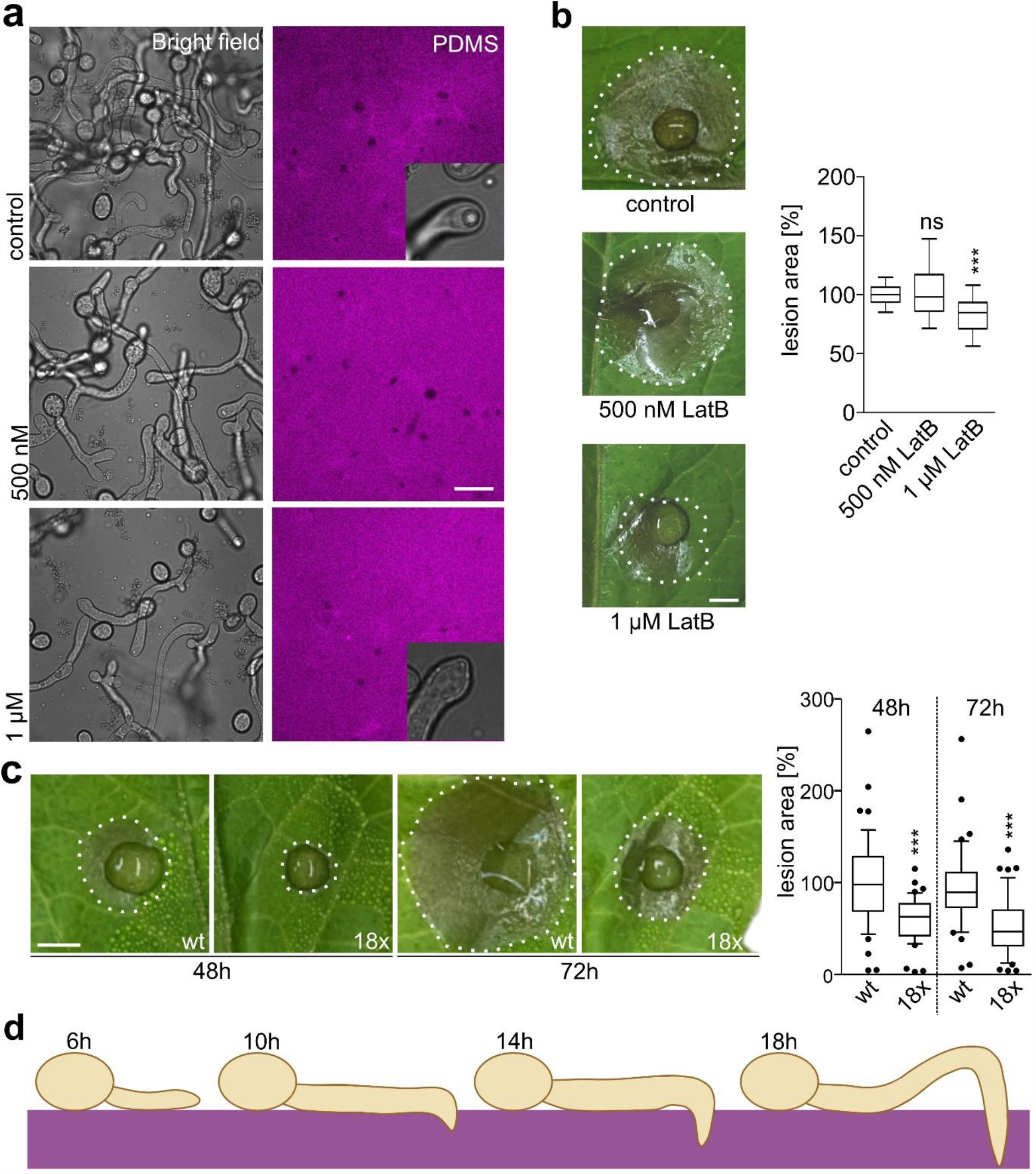
Actin filament organization is prerequisite for penetration. **(a)** Brightfield images and red fluorescence of PDMS 8 hours (18h total) after adding Latrunculin B to a final concentration of 500 nM and 1 μM. **(b)** Infection test of wildtype *Botrytis* B05.10 in presence of 0mM (n=54), 500 nM (n=27) and 1 μM (n=27) latrunculin B on tomato leaflets after 72h. Box plot showing the relative necrotic area with the wildtype mean set to 100%. Box limits in the graph represent 25th-75th percentile, the horizontal line the median and whiskers 10% to 90%. Putative differences were analyzed by One-Way ANOVA with Dunnett’s posthoc test (control: 0 mM LatB), *p<0,05; ***p<0,001, ns=not significant. **(c)** Infection test of wildtype *Botrytis* (n=48) and the 18fold (18x) mutant (n=48) on tomato leaflets after 48h and 72h. Box plot showing the relative necrotic area with the wildtype mean set to 100%. Box limits in the graph represent 25th-75th percentile, the horizontal line the median and whiskers 10% to 90%. Putative differences were analyzed by One-Way ANOVA with Dunnett’s posthoc test (control: wt), ***p<0,001. **(d)** Model of different *Botrytis* penetration stages (6h,10h, 14h, 18h). Experiments were repeated twice with similar results.

While this establishes the capacity of the necrotroph *Botrytis* to generate invasive forces, it does not answer the question if these mechanical pressures contribute to infection of plants. This prompted us to interfere directly with turgor formation during infection. To this end, we infected plants in the presence of different LatB concentrations. While 500 nM did not have a significant effect on the formation of necrotic lesions in tomato leaflets,1 μM LatB caused a significant reduction in lesion formation (Fig. 4b), indicating a role of invasive mechanical pressure for virulence. During infection *Botrytis* secretes a plethora of cutinases, cell wall degrading enzymes and cytolytic proteins to weaken preformed barriers^3^. It has been therefore assumed that necrotrohic fungi do not rely on the generation of high turgor pressure to breach into the plant cell. To elaborate on the role of turgor formation for infection, we used a multifold *Botrytis* mutant, lacking 18 important phytotoxic secreted proteins (Supplementary table 1). This mutant is based on the previously established 12fold mutant (12xbb)^4^ and includes deletion of the two most abundant polygalacturonases (PG1 and PG2) as well as the recently discovered phytotoxic proteins, SSP2^19^, Hip1^20^ and CDI1^21^. In comparison to wildtype *Botrytis*, infection of the 18fold mutant is strongly impaired in virulence as shown by significantly reduced lesions on tomato leaflets (Fig. 4c). To test whether this virulence reduction is related to invasive pressure generation, we again used fluorescently-labelled PDMS in combination with confocal microscopy to determine the generated invasive pressure. However, in comparison to the wildtype, we did not find a reduction in pressure generation in the 18fold mutant. In agreement, infection in the presence of 1 μM LatB was significantly reduced in 18fold mutant and in the wildtype (Supplementary Fig. 7), suggesting independent contributions of penetration forces and phytotoxic activity to virulence of *Botrytis*.

## Discussion

To overcome the external barriers of their hosts, necrotrophic fungi like *Botrytis* secrete a cocktail of cell wall degrading enzymes, cytolytic and phytotoxic proteins as well as different toxins^3^. Together, this induces structural weakening of preformed barriers such as cuticle, cell wall and plasma membrane. This leads to the assumption of significantly decreased mechanical plant resistance and raised the question as to what extent the generation of turgor pressure is needed for the infection process. To address this quantitatively and mechanistically, we used an artificial substrate (polydimethylsiloxane, PDMS), which mimics hydrophobicity and elasticity of plants leaves and has been used previously to investigate penetration mechanics of the oomycete *Phytophthora*^11^. The ability to neglect the phytotoxic activity of *Botrytis* renders PDMF an ideal tool to investigate turgor-driven penetration mechanics.

Our data reveal a unique and time-dependent penetration pattern by *Botrytis* on PDMS (Fig. 4d). 3D reconstructions of different stages of penetration displayed changing entry angles (Fig. 2). While initially the penetration angle was oblique as reported for the *naifu* penetration by *Phytophthora*^11^, later stages showed an increasingly perpendicular angle together with an arching of the fungal hypha above the site of penetration. This coincides with a change in adhesion pattern: whereas initially the area of adhesion is comet-shaped before the penetration site, resembling the *naifu* penetration once more. but in later stages additional adhesion zones surrounding the invasion sites were detected. The emerging adhesion pattern is a concentric isotropic adhesion area around the indentation as expected for a typical appressorium. Since zones of adherence were positive for concanavalin A, carbohydrate-binding seems to be involved in *Botrytis* surface attachment. This transition in adhesion pattern might also explain the initially observed changes in penetration angle, and together fit well with the observed unique penetration pattern.

Previously, sealing of the appressorium wall by melaninwas believed to be the prerequisite for the generation of high invasive pressures. Nevertheless, for the necrotrophic fungus *Sclerotinia sclerotiorum*, cuticle indentation has been observed^22^, supporting the necessity of mechanical force for infection. Based on electron microscopical studies, however, it was observed that the penetration peg of *Sclerotinia* had only a thin layer of cell wall which was thought to be insufficient to hold high pressure^23^. Following this evidence, it was assumed until recently that enzymatic activities were presumably more important for appressorium-mediated penetration of *Sclerotinia*^24^. Based on the contact-mechanics model described previously^11^, we calculated the invasive pressure of *Botrytis* at different stages of penetration. In accordance with the initial polarized mechanical geometry and the occurrence of a more concentric circular symmetry later, the invasive pressure was increasing from 3.7 to 5.0 bar within 8h (Figure 3). A reason for the only moderate increase in pressure might be the non-synchronized fungal penetration of PDMS. The occurrence of different stages at the same time is indicated by the relatively broad distribution of pressures at the single time points. The calculated maximal turgor pressure of >40 bar for *Botrytis* is well within the reported values for (hemi) biotrophic fungi, which form specialized appressoria^9,10^.

Our results show that for *Botrytis*, the generation of pressure and subsequent penetration is dependent on functionality of the actin cytoskeleton (Fig. 4). Recently, this has been also reported for *Phytophthora* and *Colletotrichum*^11,16,25^, indicating a conserved role of actin for appressorium development. However, pharmacological interference of actin polymerization by LatB did not only affect the *Botrytis* appressorium but directly impacted penetration as well as virulence. To differentiate between the role of turgor pressure formation and secretion of proteins for virulence, we included a 18fold *Botrytis* mutant, lacking most currently known cell wall degrading and cell-death inducing proteins, in our study. Analysis of invasive pressure as well as exogenous application of LatB displayed similar behaviour of wildtype *Botrytis* and the 18fold mutant, which indicates at least partial independency of surface penetration by brute force and facilitated host infection through secretion of phytotoxic and cell wall degrading enzymes. Taken together, our data show that the necrotrophic fungus *Botrytis* uses actin-dependent turgor generation and surface adherence to employ a unique penetration pattern to overcome host resistance.

## Materials & methods

### Cultivation of Botrytis and infection tests

*Botrytis cinerea* B05.10 was used as wildtype (wt) strain for infection tests and penetration tests on PDMS. Cultivation of *Botrytis* was performed as described previously^26^. Infection tests on tomato were carried out using 1 × 10^5^ spores per ml in Gamborg medium with 10 mM glucose. Resulting lesions were documented at 48 h and 72 h, and inoculation droplet size and lesion size were quantified (area) using the freehand tool in ImageJ. Lesion minus droplet size results in secondary lesion size which we used to calculate the ratio between compared strains. The relative infection was calculated based on B05.10 wildtype set to 100%.

### Penetration on PDMS-coated coverslips

Harvested *Botrytis* conidiospores were diluted to a concentration of 1 × 10^5^ spores per ml in Gamborg medium with 10 mM glucose. 80 μl droplets were put in the center of each coated coverslip. Polydimethylsiloxane (PDMS) was stained with perylene orange and penetration events were analyzed after 6h, 10h, 12h, 14h and 18h using a confocal laser scanning microscope. This has been done on glass simultaneously.

### Plant materials and growth conditions

For infection tests tomato plants (*S. lycopersicum*), variety “marmonde”, were grown for 4-6 weeks on soil in long-day conditions (14-h light/10-h dark) at 23°C.

### Confocal microscopy

Images were acquired using a Zeiss LSM880 AxioObserver confocal laser scanning microscope equipped with a Zeiss C-Apochromat 40×/1.2 W AutoCorr M27 water-immersion objective (INST 248/254-1). Fluorescent signals of GFP (excitation/emission 488 nm/500–571 nm) and red fluorescent signals of perylene orange (excitation/emission 594 nm/598-696 nm). For pressure measurements, z-stacks with 0.5 μm intervals and a resolution of 512 x 512 pixels were recorded. The z-stacks for 3D surface rendering were recorded with 0.2 μm and a resolution of 1024 x 1024 pixels.

### Surface deformation mapping

Surface deformation maps were reconstructed from the three-dimensional image stacks of the fluorescent PDMS following the procedure described elsewhere^11^. In brief: xy-pixels were binned in 2x2 voxels; for each voxel, the intensity profile in the z-direction was fitted to a sigmoidal function to determine the z-position of the PDMS-medium interface with sub-pixel accuracy; the resulting image was finally detilted to remove small tilt angles inherent to sample placement on the microscope table. From these data, for each invasion site, the depth *d* and radius *R*, of the indentation zone where extracted. By assuming a Hertzian indentation model for a hemisphere indenting a half-space, the Indentation force *F* is calculated with:

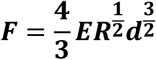

where *E* is the effective elastic modulus. For the surfaces used here, this was determined previously to be 6 bar^11^. From which in turn the maximum pressure at the invasive site is computed as:

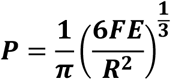

### 3D surface rendering

Surface rendering for the reconstruction of Botrytis cells and dyed PDMS was performed with Imaris 8.4 (Bitplane) software (https://imaris.oxinst.com/). For the creation of models, the surface creation tool was used for the red channel (PDMS) and the green channel (*Bcgfp1*) separately. The generated 3D models were used for angle measurements.

### Chemicals and treatments

Latrunculin B (Sigma-Aldrich, MO, USA) was used in germination/penetration tests on PDMS and added 10h after preparing the samples. For infection tests it was added to the Gamborg medium with 10 mM glucose immediately. It was obtained in powder form and dissolved in DMSO. FITC labeled concanavalin A (Sigma-Aldrich, MO, USA) was used to stain the connection between *Botrytis* hyphae and PDMS. It was obtained in powder form and dissolved in 0.9% NaCl solution.

### Generation of knockout mutants by CRISPR-Cas

Generation of multifold Botrytis gene knockout mutants was carried out as described previously^27^.

### Software

Statistical analysis was carried out using the GraphPad Prism 9 software (https://www.graphpad.com/scientific-software/prism/). The detailed statistical method employed is provided in the respective figure legends. Figures were assembled using CorelDRAW 2021. The figures 3a and 4d were generated by using the “BioRender” software.

## Supplemental data

Supplementary Fig. 1: *Botrytis* development on glass.

Supplementary Fig. 2: Occurrence of infection hyphae on PDMS.

Supplementary Fig. 3: Comparison of wildtype *Botrytis* and *bcgfp1* infection.

Supplementary Fig. 4: Fitting intensity profiles to a sigmoidal function.

Supplementary Fig. 5: Fluorescent labelling of carbohydrates by concanavalin A.

Supplementary Fig. 6: *Botrytis* germination in the presence of LatB.

Supplementary Fig. 7: Infection of wt and 18fold mutant in the presence of LatB.

Supplementary Table 1: Knockout genes of the *Botrytis* 18fold mutant.

## Acknowledgements

This work was supported by grants from the *BioComp* research initiative (Rhineland-Palatinate, Germany) and the German research foundation (DFG; SCHE 1836/5-1) to D.S.

## Contributions

T.M. performed most experiments. J.M. performed confocal microscopy. J.B. and J.S. calculated invasive and maximum pressure and provided surface deformation maps. N.S. and M.H. established the 18fold *Botrytis* knockout mutant. T.M., J.S. and D.S. designed the figures and performed statistical analysis. D.S. conceived the study and wrote the manuscript. All authors saw and commented on the manuscript.

## Corresponding author

Correspondence to David Scheuring.

## Literature

1. Hématy, K., Cherk, C. & Somerville, S. Host-pathogen warfare at the plant cell wall. Current opinion in plant biology 12, 406–413; 10.1016/j.pbi.2009.06.007 (2009).

2. Elad, Y. (ed.). Botrytis. Biology, pathology and control (Springer, Dordrecht, 2007).

3. Bi, K., Liang, Y., Mengiste, T. & Sharon, A. Killing softly: a roadmap of Botrytis cinerea pathogenicity. Trends in plant science 28, 211–222; 10.1016/j.tplants.2022.08.024 (2022).

4. Leisen, T. et al. Multiple knockout mutants reveal a high redundancy of phytotoxic compounds contributing to necrotrophic pathogenesis of Botrytis cinerea. PLoS pathogens 18, e1010367; 10.1371/journal.ppat.1010367 (2022).

5. Mendgen, K. & Deising, H. Infection structures of fungal plant pathogens - a cytological and physiological evaluation. New Phytol 124, 193–213; 10.1111/j.1469-8137.1993.tb03809.x (1993).

6. Talbot, N. J. Appressoria. Current biology : CB 29, R144–R146; 10.1016/j.cub.2018.12.050 (2019).

7. Mendgen, K. & Hahn, M. Plant infection and the establishment of fungal biotrophy. Trends in plant science 7, 352–356; 10.1016/s1360-1385(02)02297-5 (2002).

8. Money, N. P. Turgor pressure and the mechanics of fungal penetration. Can. J. Bot. 73, 96–102; 10.1139/b95-231 (1995).

9. Bechinger, C. et al. Optical measurements of invasive forces exerted by appressoria of a plant pathogenic fungus. Science (New York, N.Y.) 285, 1896–1899; 10.1126/science.285.5435.1896 (1999).

10. Loehrer, M. et al. In vivo assessment by Mach-Zehnder double-beam interferometry of the invasive force exerted by the Asian soybean rust fungus (Phakopsora pachyrhizi). New Phytol 203, 620–631; 10.1111/nph.12784 (2014).

11. Bronkhorst, J. et al. A slicing mechanism facilitates host entry by plant-pathogenic Phytophthora. Nature microbiology 6, 1000–1006; 10.1038/s41564-021-00919-7 (2021).

12. Miyoshi, M. Die Durchbohrung von Membranen durch Pilzfäden. Jahrb. Wiss. Bot. 28, 269–289 (1895).

13. Leroch, M. et al. Living colors in the gray mold pathogen Botrytis cinerea: codon-optimized genes encoding green fluorescent protein and mCherry, which exhibit bright fluorescence. Applied and Environmental Microbiology 77, 2887–2897; 10.1128/AEM.02644-10 (2011).

14. Money, N. P. Osmotic Pressure of Aqueous Polyethylene Glycols : Relationship between Molecular Weight and Vapor Pressure Deficit. Plant Physiology 91, 766–769; 10.1104/pp.91.2.766 (1989).

15. Howard, R. J., Ferrari, M. A., Roach, D. H. & Money, N. P. Penetration of hard substrates by a fungus employing enormous turgor pressures. Proceedings of the National Academy of Sciences of the United States of America 88, 11281–11284; 10.1073/pnas.88.24.11281 (1991).

16. Bronkhorst, J. et al. An actin mechanostat ensures hyphal tip sharpness in Phytophthora infestans to achieve host penetration. Science advances 8, eabo0875; 10.1126/sciadv.abo0875 (2022).

17. Coué, M., Brenner, S. L., Spector, I. & Korn, E. D. Inhibition of actin polymerization by latrunculin A. FEBS letters 213, 316–318; 10.1016/0014-5793(87)81513-2 (1987).

18. Ketelaar, T., Meijer, H. J. G., Spiekerman, M., Weide, R. & Govers, F. Effects of latrunculin B on the actin cytoskeleton and hyphal growth in Phytophthora infestans. Fungal genetics and biology : FG & B 49, 1014–1022; 10.1016/j.fgb.2012.09.008 (2012).

19. Zhu, W. et al. Botrytis cinerea BcSSP2 protein is a late infection phase, cytotoxic effector. Environmental microbiology 24, 3420–3435; 10.1111/1462-2920.15919 (2022).

20. Jeblick, T. et al. Botrytis hypersensitive response inducing protein 1 triggers noncanonical PTI to induce plant cell death. Plant Physiology 191, 125–141; 10.1093/plphys/kiac476 (2023).

21. Zhu, W. et al. Botrytis cinerea BcCDI1 protein triggers both plant cell death and immune response. Frontiers in Plant Science 14, 1136463; 10.3389/fpls.2023.1136463 (2023).

22. Lumsden, R. D. & Wergin, W. P. Scanning-Electron Microscopy of Infection of Bean by Species of Sclerotinia. Mycologia 72, 1200; 10.2307/3759575 (1980).

23. Tariq, V. N. & Jeffries, P. Ultrastructure of penetration of Phaseolus spp. by Sclerotinia sclerotiorum. Can. J. Bot. 64, 2909–2915; 10.1139/b86-384 (1986).

24. Liang, X. & Rollins, J. A. Mechanisms of Broad Host Range Necrotrophic Pathogenesis in Sclerotinia sclerotiorum. Phytopathology 108, 1128–1140; 10.1094/PHYTO-06-18-0197-RVW (2018).

25. Zhang, Y. et al. Actin-bundling protein fimbrin regulates pathogenicity via organizing Factin dynamics during appressorium development in Colletotrichum gloeosporioides. Molecular plant pathology 23, 1472–1486; 10.1111/mpp.13242 (2022).

26. Müller, N. et al. Investigations on VELVET regulatory mutants confirm the role of host tissue acidification and secretion of proteins in the pathogenesis of Botrytis cinerea. The New phytologist 219, 1062–1074; 10.1111/nph.15221 (2018).

27. Leisen, T. et al. CRISPR/Cas with ribonucleoprotein complexes and transiently selected telomere vectors allows highly efficient marker-free and multiple genome editing in Botrytis cinerea. PLoS pathogens 16, e1008326; 10.1371/journal.ppat.1008326 (2020).

