## Supplemental Data for "Plant infection by the necrotrophic fungus *Botrytis* requires actin-dependent generation of high invasive turgor pressure"

### Supplementary Figures

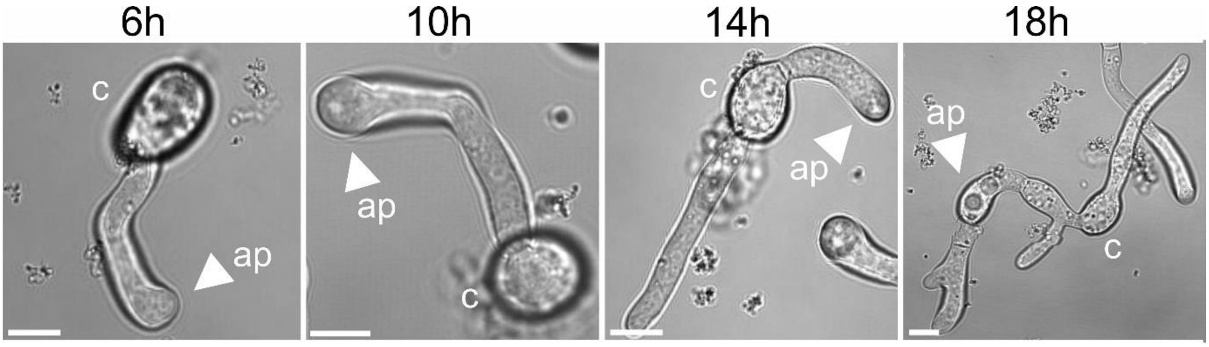


**Supplementary Fig.1: *Botrytis* development on glass.** Bright-field images of *Botrytis* conidiospores (c) germinating on glass after 6h, 10h, 14h and 18h. The appressorium (ap) is highlighted (white arrowhead). Scale bars are 5 µm.

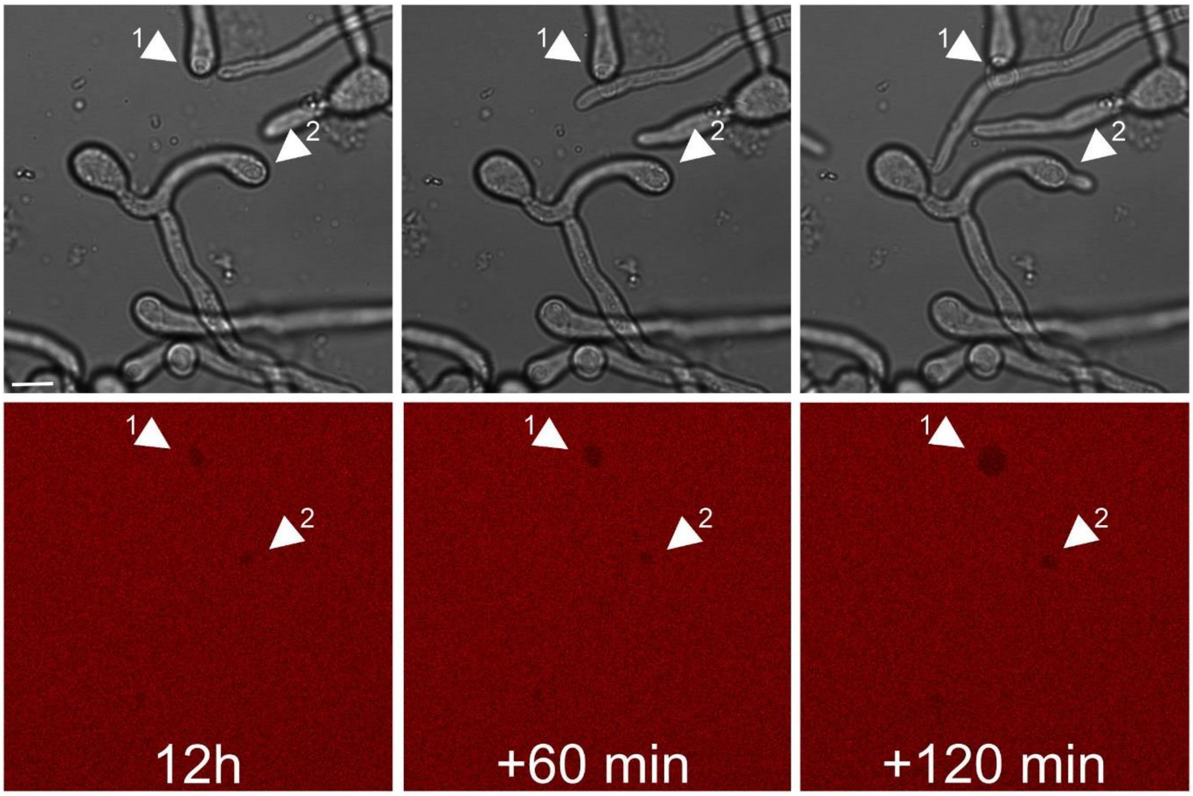


**Supplementary Fig. 2: Occurrence of infection hyphae on PDMS.** Bright-field and red fluorescence images showing *Botrytis* penetration and formation of infection hyphae on PDMS after 12h, 13h and 14h. Numbers 1 and 2 mark two separate penetration sites. Scale bar is 10 µm.

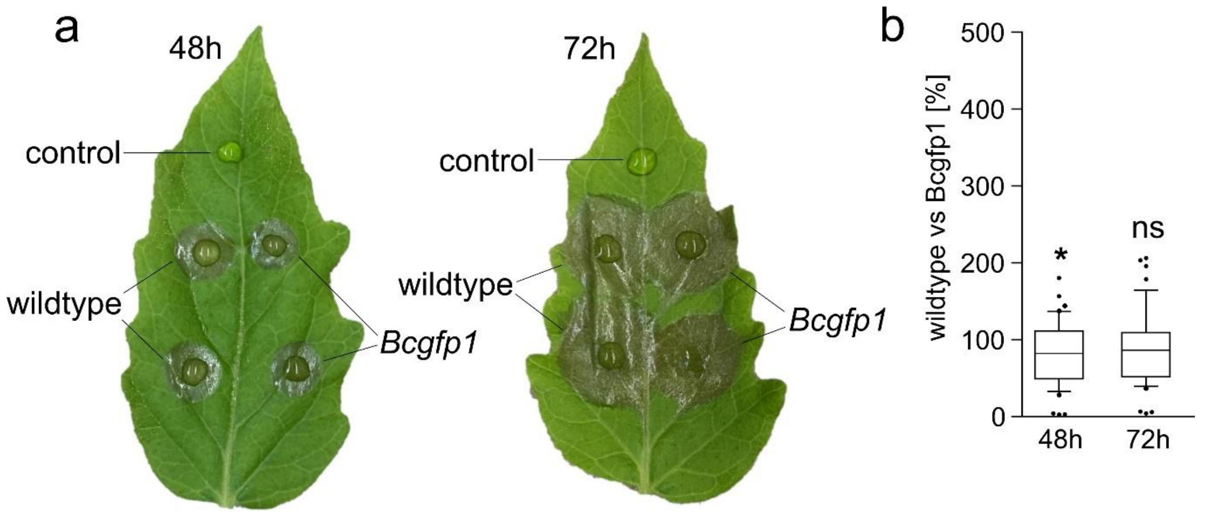


**Supplementary Fig. 3: Comparison of wildtype *Botrytis* and *Bcgfp1* infection. (a)** Infection test of wildtype Botrytis and the GFP-expressing strain *Bcgfp1* on tomato leaflets after 48h and 72h. **(b)** Box plot showing the relative necrotic area with the wildtype mean set to 100%. Box limits in the graph represent 25th-75th percentile, the horizontal line the median and whiskers 10% to 90%. Differences were analyzed by student´s t-test, *p<0,05; ns=not significant.

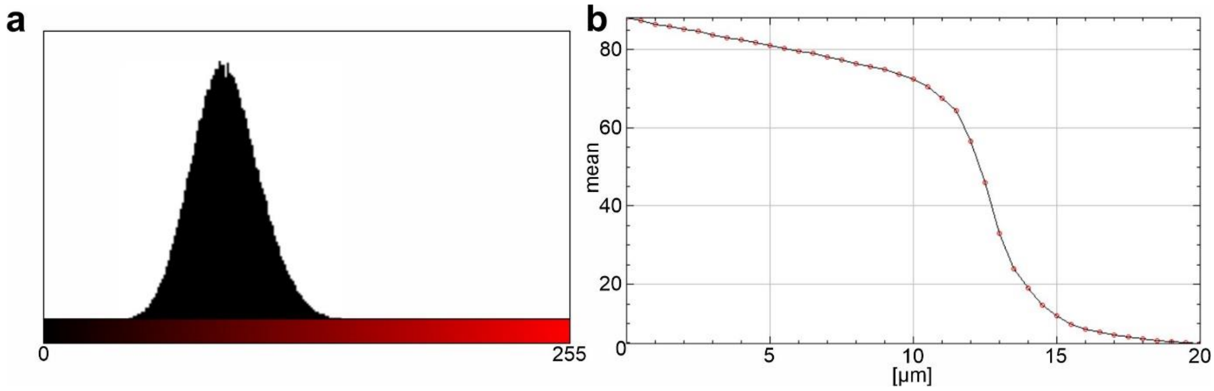


**Supplementary Fig. 4: Fitting intensity profiles to a sigmoidal function.** Example for **(a)** Gaussian distribution of fluorescence signal in the PDMS layer and **(b)** an intensity profile of a whole z stack.

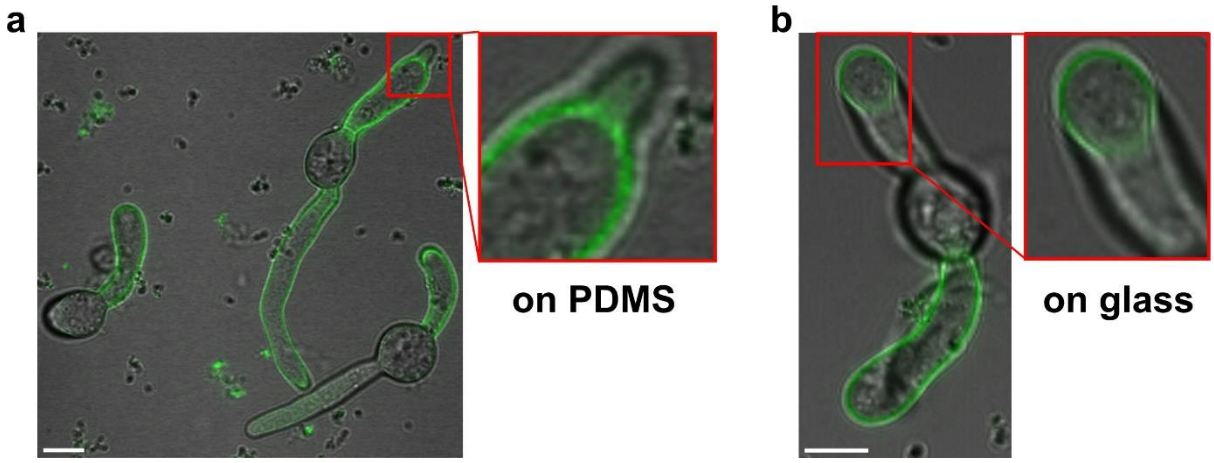


**Supplementary Fig. 5: Fluorescent labelling of carbohydrates by concanavalin A.** Bright-field images of germinating *Botrytis* conidiospores on PDMS (a) and glass (b). Germlings are stained with an Alexa fluor 488 conjugate with Concanavalin A. Scale bar is 5 µm.

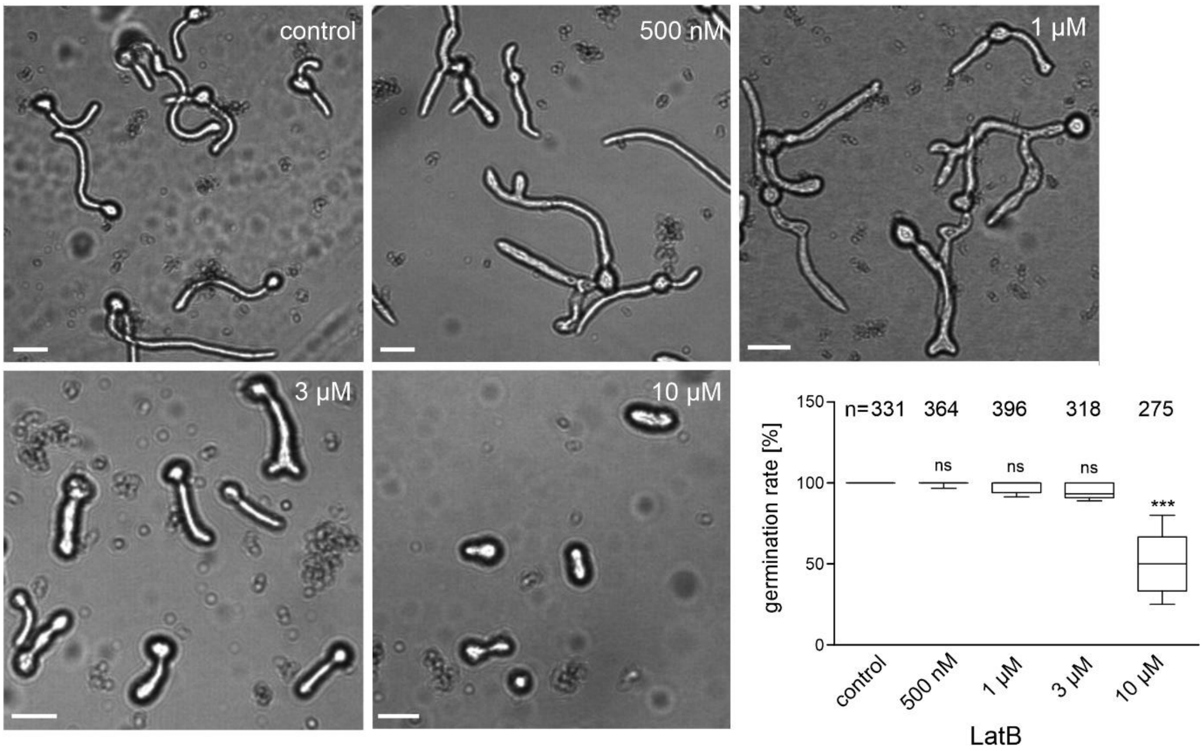


**Supplementary Fig. 6: *Botrytis* germination in the presence of Latrunculin B.** Bright-field images of *Botrytis* conidiospores germinating on a glass surface. Latrunculin B (LatB) concentrations between 500 nM and 10 µM were tested. Box plot showing the relative necrotic area with the wildtype mean set to 100%. Box limits in the graph represent 25th-75th percentile, the horizontal line the median and whiskers 10% to 90%. Putative differences were analyzed by One-Way ANOVA with Dunnett´s posthoc test (control: no LatB), ***p<0,001; ns=not significant. Scale bar is 10 µm.

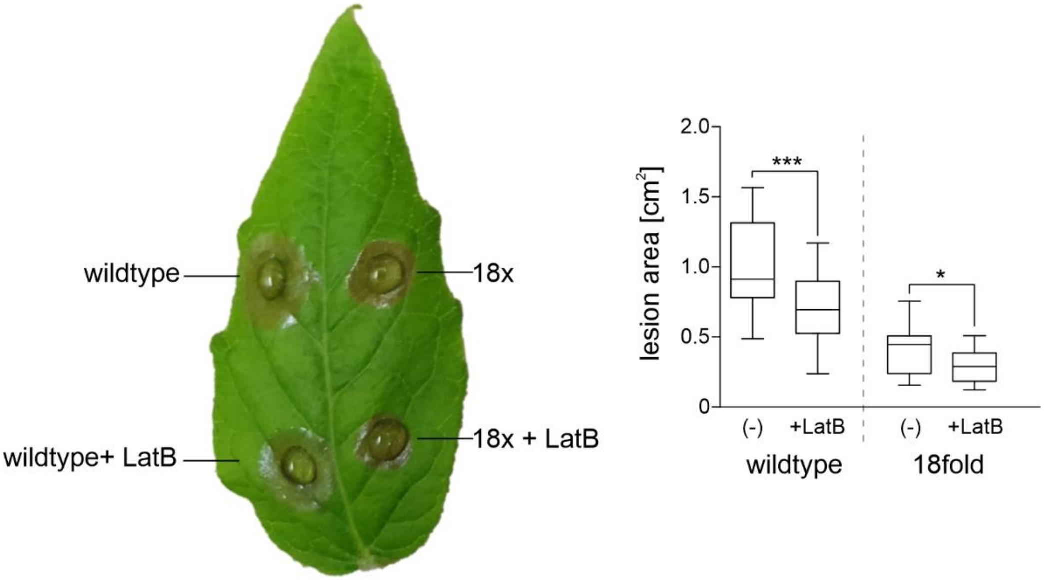


**Supplementary Fig. 7: Infection of wildtype *Botrytis* and 18fold mutant in the presence of LatB.** Lesion area was determined 72h after inoculation (n=24). Box limits in the graph represent 25th-75th percentile, the horizontal line the median and whiskers 10% to 90%. Putative differences were analyzed using by student´s t-test; *p<0,05; ***p<0.001.

**Supplementary Table 1: Gene deletions of the *Botrytis* 18fold mutant.**

| knockout order | Protein | Gene | Accession |
|---|---|---|---|
| 1 | **Spl1** | *Bcspl1* | Bcin03g00500 |
| 2 | **Nep1** | *Bcnep1* | Bcin06g06720 |
| 3 | **Nep2** | *Bcnep2* | Bcin02g07770 |
| 4 | **XYN11A** | *xyn11A* | Bcin03g00480 |
| 5 | **Hip1** | *BcHip1* | Bcin14g01200 |
| 6 | **XYG1** | *BcXYG1* | Bcin03g03630 |
| 7 | **PLP1** | *Bcplp1* | Bcin10g01020 |
| 8 | **IEB1** | *BcIEB1* | Bcin15g00100 |
| 9 | **Xyl1** | *Bcxyl1* | Bcin09g01800 |
| 10 | **GS1** | *BcGs1* | Bcin04g04190 |
| 11 | **-** | *Bcbot2* (Botrydial) | Bcin12g06390 |
| 12 | **-** | *Bcboa6* (Botcinin) | Bcin01g00060 |
| 13 | **PG1** | *Bcpg1* | Bcin14g00850 |
| 14 | **Crh1** | *Bccrh1* | Bcin01g06010 |
| 15 | **CDI1** | *BcCDI1* | Bcin06g00550 |
| 16 | **CFEM1** | *Bccfem1* | Bcin10g02180 |
| 17 | **PG2** | *Bcpg2* | Bcin14g00610 |
| 18 | **SSP2** | BcSSP2 | Bcin05g03680 |
